# Early life stress is associated with earlier emergence of permanent molars

**DOI:** 10.1101/2021.03.22.436513

**Authors:** Cassidy L. McDermott, Katherine Hilton, Anne T. Park, Ursula A. Tooley, Austin L. Boroshok, Muralidhar Mupparapu, JoAnna M. Scott, Erin E. Bumann, Allyson P. Mackey

## Abstract

Exposure to adversity can accelerate biological aging. However, existing biomarkers of early aging are either difficult to detect in individuals at scale, like epigenetic signatures, or cannot be detected until late childhood, like pubertal onset. We evaluated the hypothesis that early adversity is associated with earlier molar eruption, an easily assessed measure that has been used to track the length of childhood across primates. In a pre-registered analysis (*N* = 117, ages 4-7), we demonstrate that lower family income and exposure to adverse childhood experiences (ACEs) are significantly associated with earlier eruption of the first permanent molars, as rated in T2-weighted magnetic resonance images (MRI). We replicate relationships between income and molar eruption in a population-representative dataset (NHANES; *N* = 1,973). These findings suggest that the impact of stress on the pace of biological development is evident in early childhood, and detectable in the timing of molar eruption.

## Introduction

Exposure to early life stress, including poverty and adverse childhood experiences (ACEs), undermines physical and mental health (1). Recently, accelerated biological aging has gained attention as a potential mechanism linking experiences of poverty and maltreatment with increased risk for poor health (2). Childhood adversity has been linked to earlier puberty, faster epigenetic aging, and earlier brain maturation (3). Life history theory (4) suggests that individuals who grow up in secure, enriched environments prioritize a slower pace of development, in order to maximize parental investment and extend periods of plasticity that facilitate learning. In contrast, individuals who grow up in harsh, stressful environments may prioritize a faster pace of development, leading to earlier attainment of adult-like abilities and increased reproductive fitness, but at a steep cost to later physical and mental health.

Here, we investigate a domain of biological maturation that has not previously been considered in the childhood adversity literature: molar eruption. Molar eruption patterns have provided insights into the evolution of human development. Across broad samples of primates, age at molar eruption has been found to correlate with age at weaning, age of sexual maturity, and brain weight (5, 6, but see 7). Furthermore, the evolution of an extended childhood in humans is evident in the later age of first molar emergence and later age for completion of brain growth in humans, as compared with non-human primates and early human ancestors (8). Molar eruption is assessed routinely, unambiguously, and at an early age, as the first permanent molars typically emerge between ages 6 and 7 years. If early adversity drives early molar development, molar eruption timing could serve as an easily assessed biomarker to identify individuals at risk of accelerated development, and earlier morbidity and mortality.

## Results

We conducted a pre-registered analysis (osf.io/f6snd) to investigate how experiences of early adversity relate to the timing of emergence of the first permanent molars. We focused on two measures: family income, as low income is associated with high risk for chronic stress, and ACEs, as they have been consistently linked with poor health outcomes (e.g. 1). We measured molar eruption by rating T2-weighted MRI scans, which are sensitive to tissues in the dental follicle (Fig. 1). Our analyses included 117 four- to seven-year-old children (64 female/53 male; 48 Black/36 white/5 Asian/14 Hispanic/14 Multiracial or Other).

**Figure 1.**
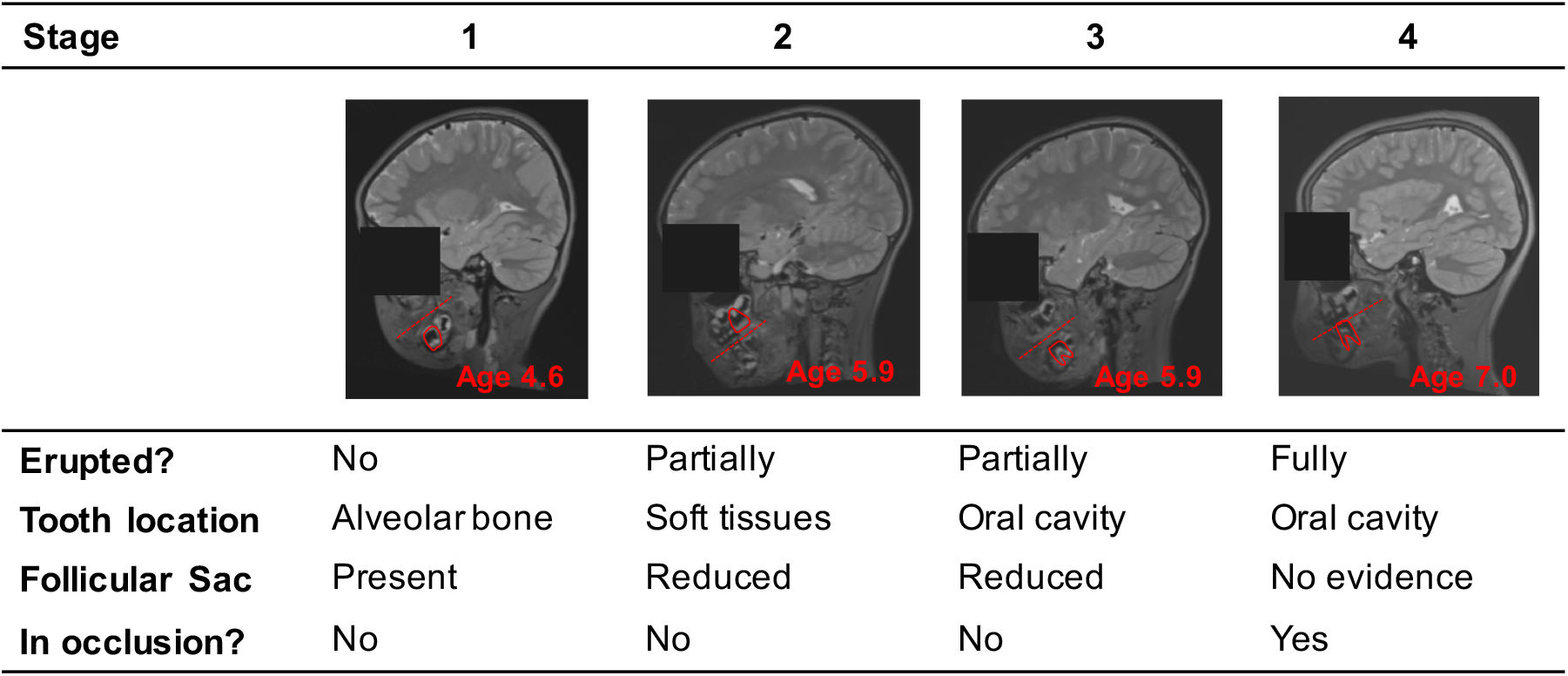
Molar eruption rating criteria in MRI.

Effects of age, gender, and race/ethnicity on molar eruption were consistent with previous reports (9). Molar eruption was positively associated with age (*β*=0.842, 95% CI [0.743, 0.942], *P*<.001; Fig. 2A). Girls had earlier molar eruption than boys (*β*=0.349, 95% CI [0.159, 0.539], *P*<.001 controlling for age; Fig. 2B). Compared with Black children, white (*β*=−0.310, 95% CI [−0.539, −0.081], *P*=.008) and Asian children (*β*=−0.599, 95% CI [−1.087, −0.111], *P*=.017) had later molar eruption. BMI was not associated with molar eruption (*β*=0.050, 95% CI [−0.063, 0.163], *P*=.381). As pre-registered, age and gender were included as covariates in all subsequent models of molar eruption, and models were run with and without race/ethnicity and BMI.

**Figure 2.**
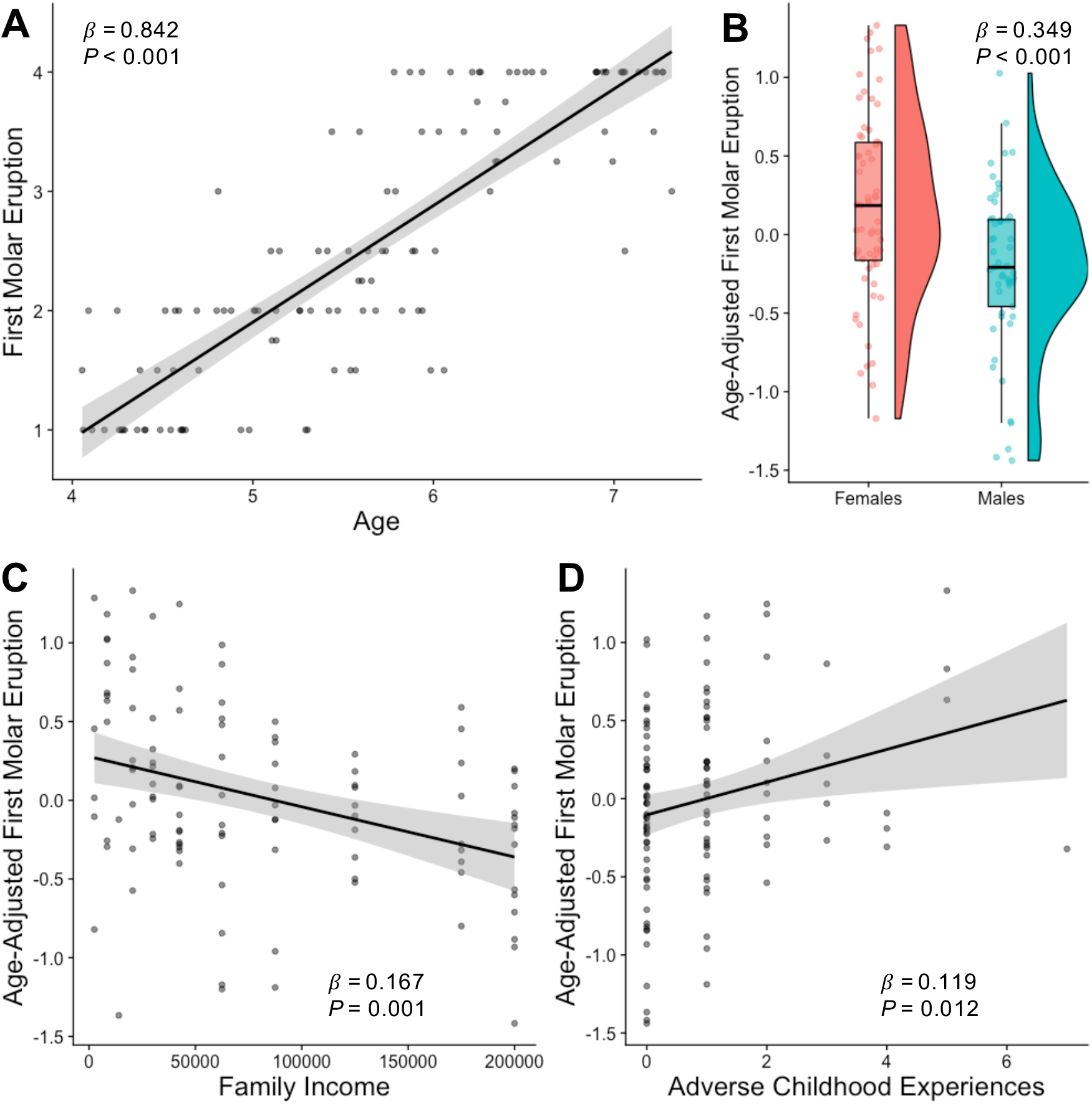
Associations between molar eruption and A) Age, B) Gender, C) Income, and D) Adverse Childhood Experiences.

Lower family income was significantly associated with earlier molar eruption (*β*=−0.167, 95% CI [−0.261, −0.073], *P*=.001 Fig. 2C). This association remained significant after controlling for BMI percentile (*β*=−0.199, 95% CI [−0.301, −0.098], *P*<.001), racial/ethnic group (*β*=−0.144, 95% CI [−0.261, −0.028], *P*=.015), or both BMI and racial/ethnic group (*β*=−0.184, 95% CI [−0.306, −0.061], *P*=.004). Greater exposure to adverse childhood experiences was also significantly associated with earlier molar eruption (*β*=0.119, 95% CI [0.027, 0.212], *P*=.012, Fig. 2D). This association remained significant when controlling for BMI (*β*=0.126, 95% CI [−0.024, 0.228], *P*=.016), but was at trend-level when including racial/ethnic group (*β*=0.086, 95% CI [−0.005, 0.178], *P*=.065). When family income and ACEs were included in the same model, only family income was significantly associated with molar eruption (income: *β*=−0.150, 95% CI [−0.246, −0.053], *P*=.003; ACEs: *β*=0.065, 95% CI [−0.030, 0.161], *P*=.178).

We conducted a parallel set of analyses using data from the National Health and Nutrition Examination Survey (NHANES; 2011-2016). Analyses were restricted to subjects between ages 4.8 and 7.8 with available oral health data, resulting in a sample of 1,973 participants. Age was related to number of first molars (*β*=0.736, 95% CI [0.712, 0.76], *P*<0.001), and girls had more first molars than boys (*β*=0.113, 95% CI [0.03, 0.196], *P*=0.01, controlling for age). Black (*β*=0.137, 95% CI [0.043, 0.23], *P*=0.006), Hispanic (*β*=0.144, 95% CI [0.072, 0.217], *P*<0.001), and multiracial children (*β*=0.156, 95% CI [0.036, 0.277], *P*=0.015) had more first molars than white children. Lower income-to-poverty ratio was significantly associated with having more first molars erupted (*β*=−0.052, 95% CI [−0.097, −0.007], *P*=0.029, controlling for age and gender). Income-to-poverty ratio was no longer significantly associated with number of first molars after controlling for BMI (*β*=−0.040, 95% CI [−0.083, 0.004], *P*=0.08) or racial/ethnic category (*β*=−0.032, 95% CI [−0.082, 0.018], *P*=0.221). Finally, we tested whether relationships between income and molar eruption extended to second molars, which erupt on average at age Among 2,993 children aged 9.1 to 14.3 years, income-to-poverty ratio was also significantly associated with number of erupted second molars (*β*=−0.044, 95% CI [−0.072, −0.017], *P*=0.003, controlling for age and gender). Income-to-poverty ratio was no longer significantly associated with number of second molars after controlling for BMI (*β*=−0.025, 95% CI [−0.053, 0.003], *P*=0.091) or race/ethnic category (*β*=−0.024, 95% CI [−0.055, 0.008], *P*=0.146).

## Discussion

Combining evidence from detailed characterization of molar eruption in MRI with nationally-representative epidemiological research, we demonstrate that children from lower income backgrounds grow up faster: their permanent molars emerge before those of their more advantaged peers. Across both analyses, Black children show earlier molar eruption than white children, which is consistent with pervasive racial disparities in income and other contextual stressors that have roots in structural racism (10). Our findings contribute to a growing literature indicating that exposure to early life stress contributes to accelerated biological maturation.

The mechanisms underlying accelerated molar eruption remain unknown. The timing of tooth emergence is regulated in part by osteocalcin, thyroid hormones, sex hormones, and cortisol (11, 12), all of which are influenced by stress (2, 13–15). We also find that controlling for BMI attenuates the relationship between income and molar eruption in NHANES, perhaps indicating that elevated BMI plays a mediating role in the effect of stress on aging processes. Work in preclinical models is needed to illuminate mechanisms linking stress to accelerated dental development.

This study has several limitations. First, we investigated family income as a contextual condition that affects stress exposure in childhood; we did not measure stress hormones or child-reported stress. Second, both samples were cross-sectional, so we cannot test for stress-induced changes in developmental trajectories. Third, the measurement of molar eruption in NHANES (present/absent) is coarse compared to techniques like MRI and X-ray, which capture variability before molars emerge into the oral cavity. Furthermore, the NHANES income-to-poverty ratio variable fails to capture differences in cost-of-living across the US that may impact the relationship between income and stress. These limitations may contribute to the weaker effect of income on molar eruption in NHANES as compared to the MRI analysis. Finally, our work was done only in the United States, and may not generalize to other contexts (16).

Despite these limitations, our work provides insight into the timing of molar eruption among children from lower-income families. Molar eruption can be characterized both in existing MRI datasets, and through routine dental care. Longitudinal research is necessary to evaluate downstream correlates of early molar eruption, including early puberty, early brain development, and mental and physical health. If molar eruption timing can identify children at risk of accelerated aging following early life stress, it may serve as a useful screening tool to direct early intervention resources to the children who need them most.

## Acknowledgments

We would like to thank all of the families who participated in this research. This study was supported by the Jacobs Foundation Early Career Award (A.P.M.), NIDA (1R34DA050297-01 to A.P.M.) and National Science Foundation Graduate Research Fellowships to C.L.M., U.A.T. and A.L.B. We thank Megan Gunnar, Ph.D., and Sara Jaffee, Ph.D. for helpful comments.

