## Supplemental Methods for "Early life stress is associated with earlier emergence of permanent molars"

#### Molar Eruption in MRI Sample

*Participants.* This study was approved by the Institutional Review Board at the University of Pennsylvania. All parents provided written informed consent. Children and their parents were recruited from the greater Philadelphia area as part of two larger studies, one on environmental impacts on brain development and learning, and one on mathematical reasoning. Participants were screened and excluded from each study if they had a diagnosed psychiatric, neurological, or learning disorder, were born more than six weeks premature, were adopted, or had any contraindications for MRI scanning.

*MRI Data Acquisition.* Imaging was performed at the Center for Advanced Magnetic Resonance Imaging and Spectroscopy (CAMRIS) at the University of Pennsylvania. Scanning was performed using a Siemens MAGNETOM Prisma 3-Tesla MRI scanner with a 32-channel head coil. A whole-brain T2-weighted sampling perfection with application optimized contrast using different angle evolutions (SPACE) structural image was acquired (acquisition parameters: TR = 3200 ms, TE = 4.06 ms, variable flip angle, voxel size = 1 mm isotropic, matrix size = 256 x 256, 176 sagittal slices, FOV = 256 mm).

*Molar eruption.* Molar eruption was rated from each subject's raw T2-weighted MRI scan by a dental student (K.H.) at the University of Pennsylvania School of Dental Medicine, in consultation with a faculty member (M.M.) with expertise in oral and maxillofacial radiology. The dental student and faculty member together developed a novel scale for classifying molar eruption status in MRI. This scale was developed after the pre-registration was submitted, but before analyses of molar eruption were performed. Eruption status of each first permanent molar was rated on a scale from 1 (unerupted) to 4 (fully erupted). The molar eruption classification can be further described as follows: In stage 1, the molar along with the follicular space is fully within the alveolar bone. In stage 2, the molar is partially erupted but submerged within the soft tissues, and the follicular space is reduced. In stage 3, the molar is partially erupted into the oral cavity but is not yet in occlusion. In stage 4, the molar is fully erupted and in occlusion (i.e. the maxillary and mandibular molars are in contact), and there is no evidence of remaining follicular space. A summary of this scale as well as example scans are shown in Fig. 1. The eruption status of all four first molars was averaged to create a single molar eruption variable that was used in analyses. While rating molar eruption, the dental student was blind to all demographic variables, including age, gender, race, ethnicity, family income, and exposure to adverse childhood experiences.

Molar eruption was rated for 133 participants under age 8. MRI scans for an additional 11 subjects were viewed by the dental student but were determined to have motion artifacts or signal dropout in the mouth that prohibited accurate rating of molar eruption. We excluded 3 subjects who had extremely delayed molar eruption (defined prior to data analysis as having Cook's Distance of greater than 4/N for the molar eruption by age regression). Finally, we restricted our analyses to include only subjects within the range for which there was variability in molar eruption. Thus, our youngest subject for molar eruption analyses was the youngest subject in the full sample who had at least one molar partially erupted (4.05 years old), and our oldest subject for molar eruption analyses was the oldest subject in the full sample with less than four fully-erupted molars (7.32 years old). After excluding 13 subjects outside of this age range, we obtained a final sample size of 117 participants with molar eruption data (64F, 55%).

*Questionnaires.* Parents reported their child's date of birth, gender, race, and ethnicity. Parents were asked to indicate each of the following race categories that applied to their child: American Indian or Alaska Native; Asian; Black or African American; Native Hawaiian or Other Pacific Islander; white; or Other. Parents were also asked to indicate whether or not their child was Hispanic or Latino. We recoded race and ethnicity into the following categories, in order to maintain consistency with NHANES race coding: Black or African American (N = 48, 41%), white (N = 36, 31%), Asian (N = 5, 4%), Hispanic (N = 14, 12%), and Multiracial or Other (N = 14, 12%).

Parents also reported their total annual family income in one of 10 income bins (less than $5000; $5000-$11,999; $12,000-$15,999; $16,000-$24,999; $25,000-$34,999; $35,000-$49,999; $50,000-$74,999; $75,000-$99,999; $100,00-$149,999; $150,000-$199,999; and $200,000 or greater). For analytical purposes, income was re-coded to represent the median of each bin, thus the maximum possible income in our sample was $200,000. Income data was available for 108 children. Parents completed the 10-item Adverse Childhood Experiences (ACE) questionnaire about their child. The ACE questionnaire is a widely-used assessment of early childhood experiences, which measures sexual, physical and emotional abuse, witnessing domestic violence, physical and emotional neglect, parental separation or divorce, and substance abuse, mental illness, or incarceration of an adult in the household (1). An ACE score is calculated by summing the responses to each of the adversity categories, with a maximum possible score of 10. Child ACEs in our sample ranged from 0 to 7, and ACEs data was available for 111 children.

*Body Mass Index.* Each child's height and weight were measured at their study visit. Height and weight data were available for 102 children. We used each child's age, gender, height and weight to calculate body mass index (BMI). We then derived BMI-by-age percentiles based on the Center for Disease Control growth charts for children and teens, using the 'childsds' package in R (2).

*Statistical Analyses.* We tested for effects of age, gender, race/ethnicity, and BMI percentile on molar eruption using linear regression. For analyses including race, the most numerous racial group in the sample (Black or African American) was treated as the reference group, and covariates were included for the other race/ethnic groups. We registered our main molar eruption analyses in an Open Science Framework pre-registration (osf.io/f6snd). First, we examined the relationship between molar eruption and childhood environment (family income and ACE score) using linear regression, with gender and age as covariates. We examined the impact of income and ACEs on molar eruption separately, and then included both environmental variables in the same model. We also ran each model with and without controlling for race/ethnicity and BMI percentile. Although our pre-registration also included re-running each model separately within each racial/ethnic group, and separately for males and females, we do not include those results here due to space limitations. We report standardized effect sizes for independent variables, obtained by running the linear models after centering and scaling all variables.

### Replication in NHANES

*Participants.* The National Health and Nutrition Examination Survey is a continuing cross-sectional survey of the civilian, noninstitutionalized US population. NHANES assesses demographic, socioeconomic, dietary, and health-related information from a sample that represents the US population of all ages. NHANES data, recommended survey methods, and analytic guidelines are available through the National Center for Health Statistics (https://www.cdc.gov/nchs/nhanes/). For this analysis, three cycles of NHANES were combined (2011-2012, 2013-2014 and 2015-2016). Analyses of first molars were restricted to include subjects with oral health data between ages 57 and 94 months, the ages for which there is variability in the number of first molars (N = 1973). Analyses of second molars were restricted to subjects aged 109 to 172 months with oral health data (N = 2993).

*Molar Eruption.* The NHANES oral health examination was conducted at the Mobile Examination Center (MEC) by licensed dentists. The number of erupted first molars was calculated from the Coronal Caries: Tooth Count variable for teeth numbers 3, 14, 19 and 30. The number of erupted second molars was calculated from the Coronal Caries: Tooth Count variable for teeth numbers 2, 15, 18, and 31. If a tooth was coded "U" (unerupted), it was considered unerupted, and otherwise the tooth was considered to be erupted. The number of erupted teeth was summed to create first and second molar eruption variables ranging from 0 to 4.

*Interview data.* Demographic information for NHANES was collected in the participants' homes by trained interviewers. Information for children under age 16 years was collected from an adult proxy in the home. Race/ethnicity was categorized as Mexican American, Other Hispanic, Non-Hispanic white, Non-Hispanic Black, Non-Hispanic Asian, or Other (including Multi-Racial). Participants could only endorse one race category. For our analyses, we combined the Mexican American and Other Hispanic groups to create a single Hispanic category. Thus, the unweighted NHANES sample composition was 23.47% white, 26.76% Black, 33.76% Hispanic, 8.77% Asian, and 7.25% Other and Multiracial. After applying sample weights (described below), weighted race/ethnicity estimates for this NHANES sample were as follows: 48.21% white, 14.95% Black, 26.43% Hispanic, 4.65% Asian, and 5.76% Other and Multiracial.

We used the NHANES income to poverty ratio (IPR) variable to estimate each participant's family income. The IPR is calculated as the ratio of household income to the poverty line based on the Department of Health and Human Services poverty guidelines. The ratio ranges from 0 (lowest income) to 5 (highest income), with values at or above 5.00 coded as 5.00.

*Body Mass Index.* Trained health technicians collected body measurements for NHANES participants during the MEC visit. We derived BMI-by-age percentiles, as described above.

*Statistical Analyses.* Analyses were completed using the R package 'survey' version 4.0 (3). New sample weights for the 6-year sample were calculated by dividing the 2-year sample weights by 3, as recommended in NHANES documentation. In order to replicate our analyses in the MRI sample, we first examined the relationship between number of first molars and family income using linear regression, controlling for age and gender. We then repeated the model controlling for race/ethnicity and BMI percentile. We report standardized effect sizes for independent variables in NHANES analyses.
